# New Dual Inducible Cellular Model to Investigate Temporal Control of Oncogenic Cooperating Genes

**DOI:** 10.1101/2024.02.23.581802

**Authors:** Matthew R. Kent, Amanda N. Jay, Genevieve C. Kendall

## Abstract

The study of cooperating genes in cancer can lead to mechanistic understanding and identifying potential therapeutic targets. To facilitate these types of studies, we developed a new dual-inducible system utilizing the tetracycline- and cumate-inducible systems driving HES3 and the PAX3::FOXO1 fusion-oncogene, respectively, as cooperating genes from fusion-positive rhabdomyosarcoma. With this new model, we can independently induce expression of either HES3 or PAX3::FOXO1, as well as simultaneously induce expression of both genes. This new model will allow us to further investigate the cooperation between HES3 and PAX3::FOXO1 including the temporal requirements for genetic cooperation. This dual-inducible model can be adapted for any cooperating genes, allowing for independent, simultaneous, or temporally controlled gene expression.

## Introduction

Functional genomics is an important strategy for understanding oncogenesis and therapeutic vulnerabilities. Loss and gain-of-function approaches in a cell culture context can be leveraged to define genetic cooperation and the resulting phenotypic outcomes. To this end, overexpression or knockdown of putative oncogenes and cooperating genes has led to important discoveries in how they may be functioning in cancer cells [1-5]. However, typically, this type of modeling does not account for critical factors in the tumorigenic process, including timing of expression, the level and duration of expression, or cessation of expression. For these types of questions, an inducible system such as the tetracycline-inducible system or the cumate-inducible system is much better suited. Additionally, combining these systems to have temporal control over multiple genetic events is challenging because of the required number of vectors to introduce into cells.

The tetracycline-inducible system utilizes either tetracycline or a derivative, such as doxycycline, to reversibly induce transcriptional activation or repression [6]. In a Tet-On system, a combination of a tetracycline-controlled transcriptional silencer (tTS), which silences TRE promoters in the absence of tetracycline, and a reverse tetracycline transactivator (rtTA), which is only capable of binding the TRE in the presence of tetracycline, are used (Gossen et al, 1995. Science). The cumate repressor system utilizes the bacterial cumate repressor (CymR), which binds to the cumate operon (CuO) in the absence of cumate [7]. The tetracycline-inducible and cumate-inducible systems are each meant to be used with a single gene at a time given the packaging limitations for vectors backbones, thus these systems are challenging to use for complex studies of gene cooperation.

In this study we combined these two systems in a single cell line via a series of vectors introduced by lentiviral transduction. We engineered the regulatory tTS and rtTA for the Tet-On system and the cumate repressor CymR, all separated by viral T2A sequences, to be in one vector. In a second vector, we designed a tetracycline-inducible HES3 tagged with EGFP by a viral T2A sequence. In the third vector, we designed a cumate-inducible PAX3::FOXO1 tagged with mCherry by a viral T2A sequence. By utilizing both the Tet-On and cumate repressor systems in a single cell line, we generated a novel system to investigate potential cooperation of two oncogenes, independently controlling both the level and timing of expression of each gene.

## Materials and Methods

### Cell culture

293T cells (CRL-3216, ATCC, RRID: CVCL_0063) were grown in Dulbecco’s modified Eagle’s media with GlutaMAX (DMEM, 10569044, Gibco) supplemented with 10% FBS, 1x Penicillin/Streptomycin, and 10 mM glutamine. Cells were passaged every 3-4 days with TrypLE (12604013, Gibco). Cells were authenticated by STR and tested for mycoplasma annually through Genetica Inc a subdivision of LabCorp.

### Lentiviral plasmids

To design the plasmid containing the Tet-On regulatory proteins and the cumate repressor, we used VectorBuilder’s Tet Regulatory Protein Expression Lentiviral Vector as a base, which contains both the tTS and rtTA proteins linked via a T2A sequence. We then added the CymR sequence directly downstream of this linked by a second T2A sequence. Additionally, this plasmid contains a blasticidin resistance gene for antibiotic selection. The plasmid map is available in Supplemental Figure S1.

To design the plasmid containing the tetracycline-inducible gene, we used VectorBuilder’s Mammalian Tet Inducible Gene Expression Lentiviral Vector as a base, adding the coding sequence for either a multicloning site (MCS) or human HES3 with no stop codon directly downstream of the TRE sequence, linked to EGFP with a T2A sequence. Additionally, this plasmid contains a puromycin resistance gene for antibiotic selection. The plasmid maps are available in Supplemental Figure S2 for MCS version and Supplemental Figure S3 for HES3 version.

To design the plasmid containing the cumate-inducible gene, we used VectorBuilder’s Mammalian Tet Inducible Gene Expression Lentiviral Vector as a base. Then we replaced the TRE sequence with a CMV promoter and CuO sequence, adding the coding sequence for either a multicloning site (MCS) or human PAX3::FOXO1 with no stop codon directly downstream of the TRE sequence, linked to mCherry with a T2A sequence. Additionally, this plasmid contains a hygromycin resistance gene for antibiotic selection. The plasmid maps are available in Supplemental Figure S4 for MCS version and Supplemental Figure S5 for PAX3::FOXO1 version.

These plasmids were generated and packaged into lentivirus by VectorBuilder with a minimum virus titer of 1×10^8^ TU/mL.

### Lentiviral transduction

293T cells were transduced with each vector sequentially to prevent potential cytotoxicity from PAX3::FOXO1 expression. The multiplicity of infection (MOI) used for each vector was 5. For transduction, 10,000 cells were plated on a 6-well plate with 2mL of culturing media and allowed to adhere for 16 hours overnight. Then, transduction media was made with 80% culturing media, 20% TransDux MAX Lentivirus Transduction Enhancer (LV860A-1, System Biosciences), and 0.5% TransDux (LV860A-1, System Biosciences). 72 hours after transduction, transduction media was removed and fresh culturing media was added with selection antibiotic. For the tTS/rtTA/CymR vector (developed in this study, Supplemental Figure S1), 3µg/mL of blasticidin (A1113903, Fisher Scientific) was used for selection. For the TRE:HES3-T2A-EGFP vector (developed in this study, Supplemental Figure S3), 1µg/mL of puromycin (A1113803, Fisher Scientific) was used for selection. For the CMV-CuO:PAX3::FOXO1-T2A-mCherry vector (developed in this study, Supplemental Figure S5), 300µg/mL of hygromycin (10-687-010, Fisher Scientific) was used for selection. These concentrations were determined by antibiotic selection kill-curve. Briefly, 10,000 cells were plated in each well of a 96-well plate in growth media. In triplicate, titrated concentrations of blasticidin, puromycin, or hygromycin were added to each well every two days for six days total. After six days of exposure to antibiotic, crystal violet staining was done. After aspirating out all growth media, the cells were fixed with 4% paraformaldehyde (50-276-31, Fisher Scientific) for 15 minutes at room temperature and then washed with 1x PBS. After aspirating out the PBS, crystal violet stain was added for 5 minutes (250mg crystal violet (C0775, Sigma-Aldrich) added to 100mL of 20% methanol). The crystal violet stain was then removed, and the plate was gently washed by submerging in a container of still tap water, and repeated once with fresh tap water. The plate was then allowed to air dry. To extract and quantify, 10% glacial acetic acid in water was added to each well and vigorously agitated for 30 seconds. The plate was then read on a plate reader at 590nm.

### Tetracycline and Cumate induction

Dual-inducible 293T cells that were successfully transduced with all three vectors were plated on either 6-well or 12-well plates and allowed to adhere for 16 hours overnight. Anhydrotetracycline (C4291, ApexBio Technology) was used as an effector for the Tet-On system and binds the rtTA protein at a much higher affinity than tetracycline [8]. Anydrotetracycline was dissolved in dimethyl sulfoxide (DMSO, D2650, Sigma-Aldrich) and added in concentrations ranging from 1ng/mL to 50ng/mL. The resulting DMSO concentrations were kept to a maximum of 0.1% total volume. A water-soluble cumate solution (QM150A-1, System Biosciences) was used to induce PAX3::FOXO1-T2A-mCherry, and was added in concentrations ranging from 1µg/mL to 50µg/mL.

### Western Blotting

Cells used for western blotting were collected with TrypLE, spun down at 8000xg for 5 minutes, aspirated of supernatant media, and snap frozen to be stored at -80°C. Cells were then lysed with RIPA buffer and 1x protease inhibitor (PI78442, Fisher Scientific) for 2 hours on ice. Protein concentration was determined by BCA assay (PI23227, Fisher Scientific). For each sample, 20µg of protein was loaded into a 4-15% gradient mini-PROTEAN TGX precast protein gel (4561086, Bio-Rad). Gels were run at 150V in 1x Tris/Glycine/SDS buffer (1610772, Bio-Rad) until the dye front has just run off the gel, approximately 45 minutes. The samples were then transferred to a PVDF membrane at 400mA for 2 hours at 4°C in 1x Tris/Glycine buffer (1610771, Bio-Rad). Membranes were then blocked in casein blocking buffer (PI37528, Fisher Scientific) with 0.05% Tween-20 for 1 hour at room temperature. Membranes were then incubated overnight at 4°C with primary antibodies in fresh casein blocking buffer with 0.05% Tween-20. Primary antibodies used were mouse anti-alpha Tubulin (3873S, Cell Signaling Technologies, RRID: AB_1904178) at 1:1000, mouse anti-HES3 (PCRP-HES3-1A10, Developmental Studies Hybridoma Bank, RRID: AB_2618684) at 0.5µg/mL, and rabbit anti-FOXO1 (2880S, Cell Signaling Technologies, RRID: AB_2106495) at 1:1000. Then membranes were washed three times for 10 minutes each in 1x phosphate buffered saline (PBS) with 0.05% Tween-20. Membranes were then incubated at room temperature for 2 hours with secondary antibodies in fresh casein blocking buffer with 0.05% Tween-20. Secondary antibodies used were goat anti-mouse horse radish peroxidase (HRP)-conjugated (1706516, Bio-Rad, RRID: AB_11125547) at 1:10,000 and goat anti-rabbit HRP-conjugated (1721019, Bio-Rad, RRID: AB_11125143) at 1:10,000. Membranes were then washed three more times for 10 minutes each in 1x PBS with 0.05% Tween-20. Membranes were then imaged on a C-DiGit Chemiluminescent Western Blot Scanner (103375-240, VWR) using SuperSignal West Pico PLUS Chemiluminescent Substrate (PI34577, Fisher Scientific) to image TUBULIN and PAX3::FOXO1/FOXO1, and SuperSignal West Atto Ultimate Sensitivity Substrate (38554, Fisher Scientific) to image HES3.

### Imaging

Images were taken with a Leica DMIL LED microscope with a 10x objective. Filters for brightfield, GFP, and RFP/mCherry were used.

### Fluorescent-activated cell sorting

Live cell sorting was done using the BD Influx Cell Sorter (BD Biosciences) by BD FACS Sortware version 1.2.0.142. Cells were sorted into DMEM GlutaMAX growth media supplemented with 10% FBS, 1x Penicillin/Streptomycin, and 10mM glutamine. The sorted cell population was generated by sorting dual-induced cells for both GFP and mCherry fluorescence. The sorted cells were then plated on a T-175 flask in additional growth media with 3µg/mL blasticidin, 1µg/mL puromycin, and 300µg/mL hygromycin. Cells that were transduced and unsorted were used in Figures 1 and 2, and cells that were sorted were used in Figure 3. Both populations were able to generate reproducible induction of HES3 and PAX3::FOXO1.

**Figure 1.**
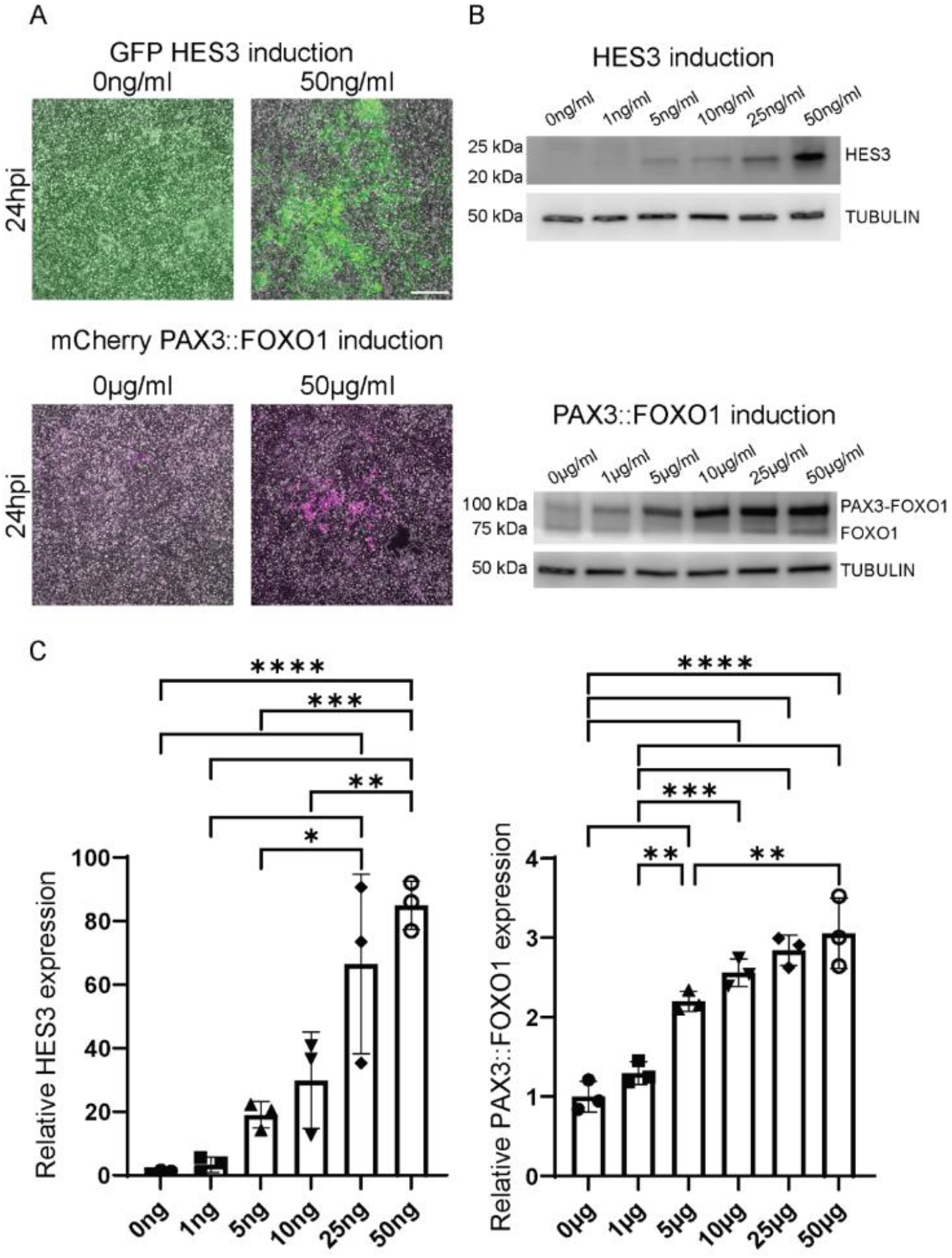
Anhydrotetracycline induction of HES3 and cumate induction of PAX3::FOXO1 is titratable. 293Tii cells were plated on 6-well plates at a density of 250,000 cells per well. After overnight adherence, cells were treated with titrating amounts of either: anhydrotetracycline at 0, 1, 5, 10, 25, or 50 ng/ml; or cumate at 0, 1, 5, 10, 25, or 50 µg/ml. Shown are representative images of cells imaged at 24 hours post-induction (hpi) on a Leica DMI microscope with a 10x objective for GFP fluorescence (HES3) or mCherry fluorescence (PAX3::FOXO1). Scale bar is 500 µm. (**A**,**B**) and then harvested for western blotting. Cells were lysed in RIPA buffer, and then 20µg of protein was loaded into each well of a 4-15% gradient gel. After transferring to a PVDF membrane, the membrane pieces were blotted with either a HES3 antibody (**C**), FOXO1 antibody that recognizes the PAX3::FOXO1 fusion and endogenous FOXO1 (**D**), or TUBULIN antibody (**C**,**D**). Each point represents a biological replicate (n=3 for each condition). The error bars represent the mean ± standard deviation. The p values were calculated using a one-way ANOVA followed by Tukey’s multiple comparisons post hoc test. This was repeated three times. Anhydrotetracycline p-values: 0ng vs 25ng, p=0.00081; 0ng vs 50ng, p=0.00008; 1ng vs 25ng, p=0.00112; 1ng vs 50ng, p=0.0001; 5ng vs 25ng, p=0.0108; 5ng vs 50ng, p=0.00075; 10ng vs 50ng, 0.00348. Cumate p-values: 0µg vs 5µg, p=0.0005; 0µg vs 10µg, p=0.00004; 0µg vs 25µg, p=0.00001; 0µg vs 50µg, p=0.000002; 1µg vs 5µg, p=0.0055; 1µg vs 10µg, p=0.0003; 1µg vs 25µg, p=0.00004; 1µg vs 50µg, p=0.00001; 5µg vs 50µg, p=0.008.

**Figure 2.**
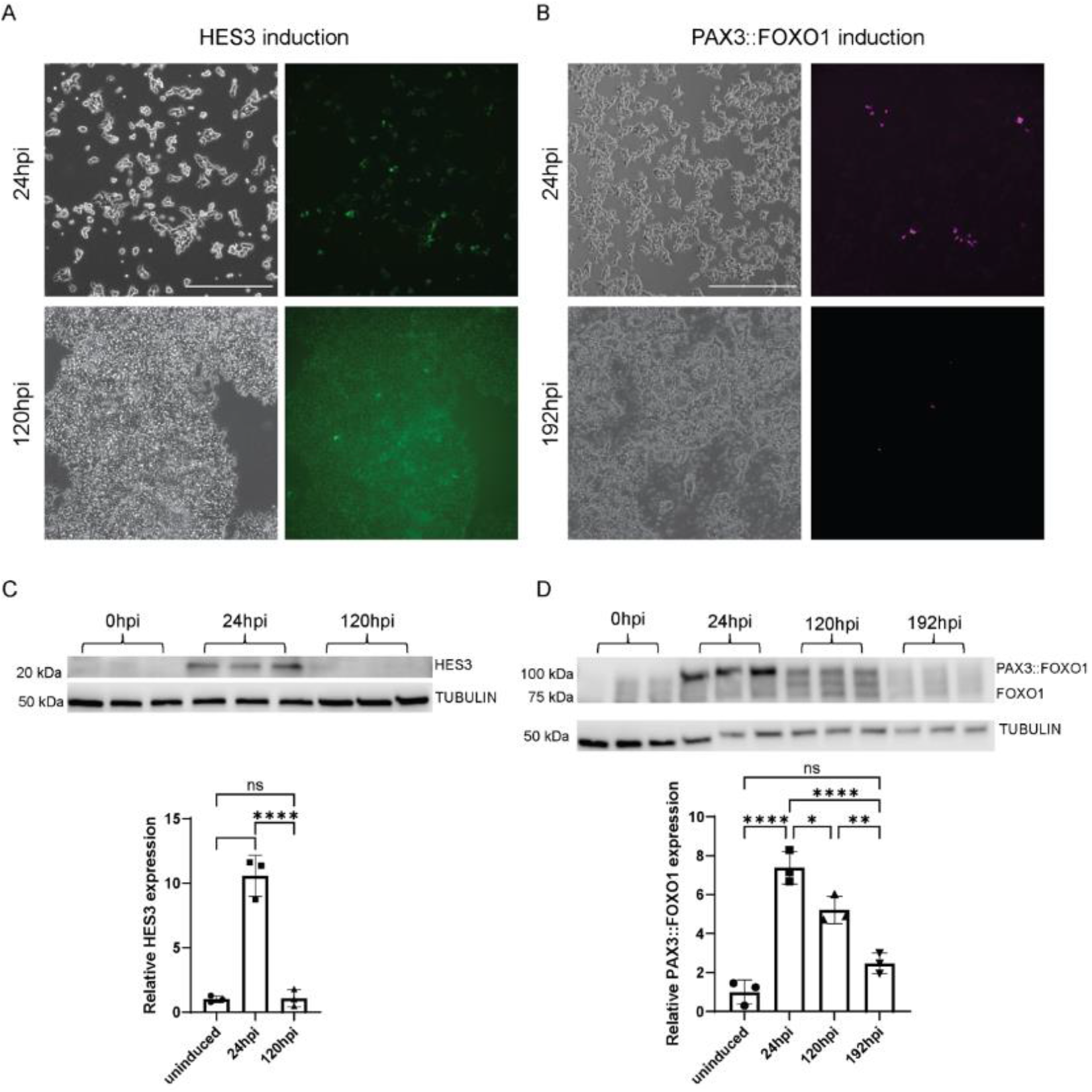
HES3 and PAX3::FOXO1 induction are both reversible. 293Tii cells were plated on 6-well plates at a density of 250,000 cells per well. After overnight adherence, cells were treated with either 50ng/ml anhydrotetracycline (HES3 induction) or 50µg/ml cumate (PAX3::FOXO1 induction). Shown are representative images of anhydrotetracycline-induced cells at 24 hours post-induction (hpi) and 120hpi (**A**), and cumate-induced cells at 24hpi, 120hpi, and 192hpi (**B**). Cells were imaged on a Leica DMI microscope with a 10x objective. Scale bar is 500 µm. Cells were then harvested for western blotting. Cells were lysed in RIPA buffer, and 20µg of protein was loaded into each well of a 4-15% gradient gel. After transferring to a PVDF membrane, the membrane pieces were blotted with either a HES3 antibody (**C**), FOXO1 antibody that recognizes the PAX3::FOXO1 fusion and endogenous FOXO1 (**D**), or TUBULIN antibody (**C**,**D**). Relative expression of either HES3 (**C**) or PAX3::FOXO1 (**D**) was then quantified using ImageJ (relative to the uninduced cells). Each point represents a biological replicate (n=3 for each condition). The error bars represent the mean ± standard deviation. The p values were calculated using a one-way ANOVA followed by Tukey’s multiple comparisons post hoc test. This was repeated three times. HES3 reversibility p-values: uninduced vs 24hpi, p=0.00006; 24hpi vs 120hpi, p=0.00006; uninduced vs 120hpi, p=0.99. PAX3::FOXO1 reversibility p-values: uninduced vs 24hpi, p=0.00001; uninduced vs 120hpi, p=0.0003; uninduced vs 192hpi, p=0.11; 24hpi vs 120hpi, p=0.019; 24hpi s 192hpi, p=0.00009; 120hpi vs 192hpi, p=0.005.

**Figure 3.**
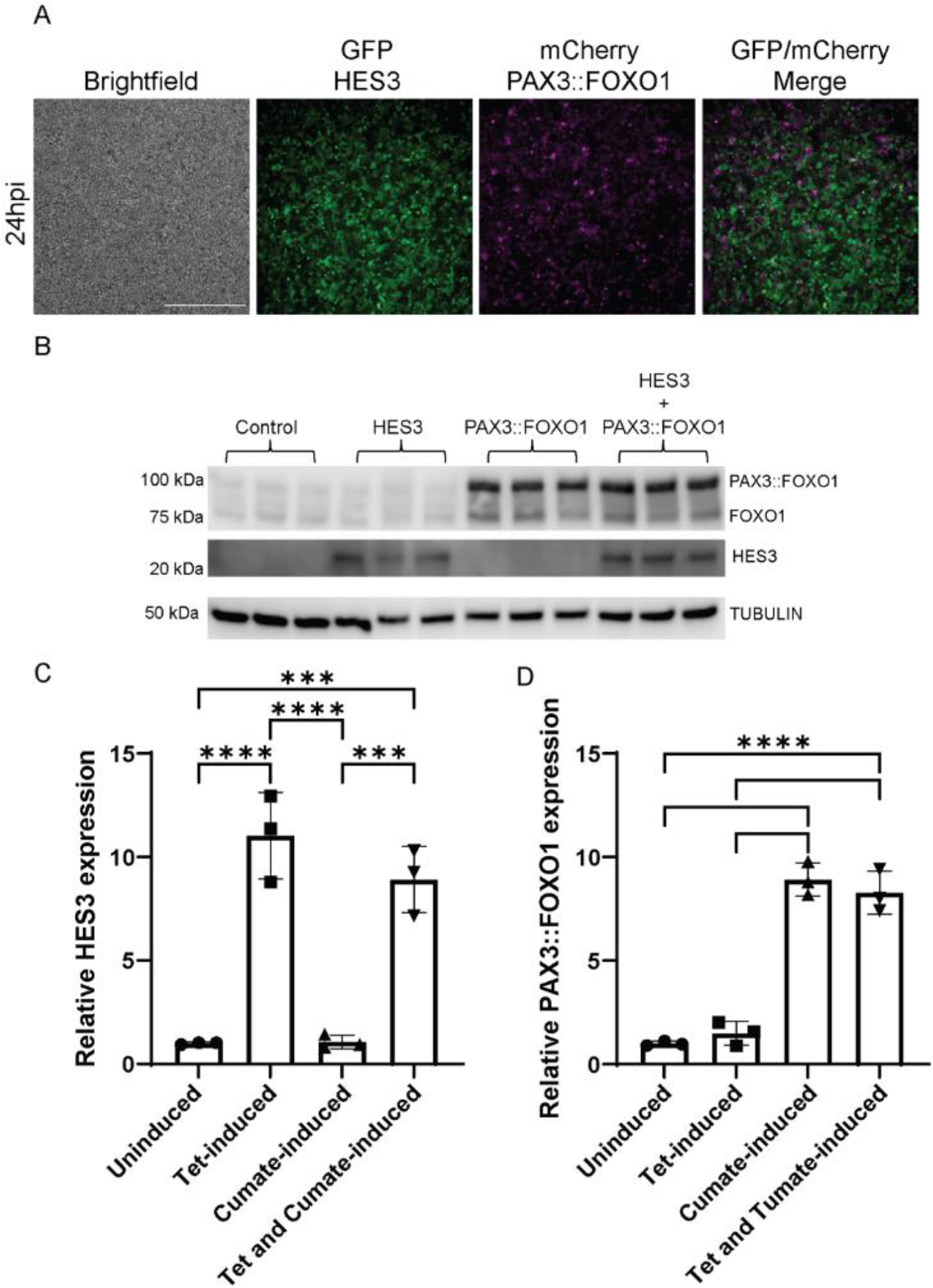
HES3 and PAX3::FOXO1 can be independently or simultaneously induced in post-sort 293Tii cells. Post-sort 293Tii cells were plated on 6-well plates. After overnight adherence, cells were treated with either 50ng/ml anhydrotetracycline (**A**), 50µg/ml cumate (**B**), or both 50ng/ml anydrotetracycline and 50µg/ml cumate (**C**). Cells were imaged at 24 hours post-induction (hpi) (**A-C**) Cells were imaged on a Leica DMI microscope with a 10x objective. Scale bar is 500 µm. Treated cells were harvested for a western blot and were lysed in RIPA buffer, and then 20µg of protein was loaded into each well of a 4-15% gradient gel. After transferring to a PVDF membrane, the membrane pieces were blotted with either a HES3 antibody, FOXO1 antibody, or TUBULIN antibody (**D**). Relative expression of HES3 and PAX3::FOXO1 was then quantified. Each point represents a biological replicate (n=3 for each condition). The error bars represent the mean ± standard deviation. The p values were calculated using a one-way ANOVA followed by Tukey’s multiple comparisons post hoc test. This was repeated three times. HES3 expression p-values: uninduced vs tet-induced, p=0.00007; uninduced vs cumate-induced, p=0.9999; uninduced vs tet- and cumate-induced, p=0.0004; tet-induced vs cumate-induced, p=0.00007; tet-induced vs tet- and cumate-induced, p=0.28; cumate-induced vs tet- and cumate-induced, p=0.0004. PAX3::FOXO1 expression p-values: uninduced vs cumate-induced, p=0.000004; uninduced vs tet- and cumate-induced, p=0.000007; tet-induced vs cumate-induced, p=0.000006; tet-induced vs tet- and cumate-induced, p=0.00001.

### Image Quantification and Statistics

Western blots were quantified using ImageJ version 1.53h (RRID: SCR_003070). Each experiment was repeated 3 times independently, and the number of samples are noted in each figure legend. Statistics were performed using GraphPad Prism 9 (RRID: SCR_002798). p-values for western blot quantifications were calculated using a one-way ANOVA followed by Tukey’s multiple comparisons post hoc test.

## Results

To generate a dual-inducible cell line, we utilized lentiviruses to transduce HEK293T cells with a series of three different constructs with the Tet-On and Cumate-On systems. The first construct contained both the reverse tet transactivator and CymR separated by a T2A sequence (Supplemental Fig S1). The second construct contained our first Gene of Interest (GOI) cDNA, HES3, separated from an EGFP reporter by a viral T2A sequence and under control of the tet response element (Supplemental Fig S3). The third construct contained our second GOI, PAX3::FOXO1, separated from an mCherry reporter by a T2A sequence and under control of the cumate response element (Supplemental Fig S5). Each construct also contained an antibiotic selection marker: blasticidin, hygromycin, or puromycin, respectively.

After transducing all three constructs, we tested that expression of HES3 and PAX3::FOXO1 could be induced by anhydrotetracycline or cumate, respectively. For HES3 induction, we titrated anhydrotetracycline, an analog of tetracycline that binds the reverse tet transactivator, at 0 ng/ml, 1 ng/ml, 5 ng/ml, 10 ng/ml, 25 ng/ml, and 50 ng/ml. For PAX3::FOXO1 induction, we titrated cumate at 0 µg/ml, 1 µg/ml, 5 µg/ml, 10 µg/ml, 25 µg/ml, and 50 µg/ml. At 24 hours post induction (hpi), we were able to see induction of EGFP (Fig 1A) and mCherry (Fig 1B). Cells were also collected for western blotting for HES3 or PAX3::FOXO1 at 24 hours post induction (hpi). We observed that an increase in drug concentration was consistent with an increase in HES3 (Fig 1C) and PAX3::FOXO1 (Fig 1D) protein expression.

We next wanted to ensure that induction of HES3 and PAX3::FOXO1 expression were reversible. To do this, we induced HES3 and PAX3::FOXO1 expression in separate cell populations. At 24hpi, we imaged the cells and harvested a portion to confirm protein induction by western blot (Fig 2A-B). Then, we replaced the cell growth media with fresh, drug-free media. From here, we imaged the cells at 48hpi, 96hpi, and 120hpi. By 120hpi, there was virtually no EGFP expression visible, and mCherry expression had decreased as well (Fig 2A-B). At 120hpi, while there was no HES3 protein expression detectable by western blot (Fig 2C), there was still detectable PAX3::FOXO1 (Fig 2D). However, by 196hpi, PAX3::FOXO1 expression decreased back to baseline expression levels. This suggests that PAX3::FOXO1 has a relatively longer half-life than HES3, but that both systems are reversibly controllable.

While we have confirmed induction of HES3 and PAX3::FOXO1 by fluorescent imaging and protein expression, we wanted to ensure that our cell population was enriched for inducible cells. To enrich our cell population for those that express our vectors, we plated one million cells and induced with anhydrotetracycline and Cumate on the same cell population for 48 hours, along with individually induced cell populations. We then harvested these cell populations and performed live fluorescence-activated cell sorting, with 29.81% of singlets being both EGFP- and mCherry-positive (Fig S6). These EGFP and mCherry dual positive cells were collected and expanded in culture. This dual induction of HES3 and PAX3::FOXO1 was further validated via imaging and western blot using the post-sorted cells. For this, we had four conditions of treated cells: water and DMSO only; anhydrotetracycline and water for HES3 induction; cumate and DMSO for PAX3::FOXO1 induction; and anhydrotetracycline and cumate for dual HES3 and PAX3::FOXO1 induction. At 24hpi, we saw the expected individual HES3 (Fig 3A) and PAX3::FOXO1 (Fig 3B) expression. We also observed simultaneous induction of HES3 and PAX3::FOXO1 (Fig 3C). This was further confirmed by western blot, in which the only samples that displayed strong expression of both HES3 and PAX3::FOXO1 were those that received both anhydrotetracycline and cumate, indicating the discrete control that is available with this system for studying cooperating oncogenes (Fig 3D).

## Discussion

Here, we demonstrated a novel system of simultaneously utilizing both the Tet-On system and the cumate repressor system in a cell line transduced with three vector constructs. We confirmed that both the Tet-on and cumate systems result in titratable levels of each gene of interest (Fig 1), and that the expression of both genes is reversible (Fig 2). Additionally, the systems are simultaneously inducible allowing for controllable expression of both genes of interest in the same cells (Fig 3).

Previous efforts to generate dual-inducible cell lines were typically done via transient transfection of plasmid DNA, allowing for only short-term study [9,10]. The system described here, similar to the PiggyBac system recently published [11], integrates the lentiviral vectors into the cell genome to generate a stable cell line rather than a transient one. This allows for more long-term and complex studies investigating potential relationship and cooperation between the two genes of interest. However, while the PiggyBac system which utilizes transfection to incorporate plasmids into the genome, has trouble integrating into non-dividing and primary cells, lentivirus does not share these limitations [12]. Therefore, our developed system could be more universally applicable to cells that are traditionally challenging to transfect.

Unlike standard overexpression or knockdown studies, this system can be used to investigate timing-dependent relationships between two genes. Additionally, it can be used to look at dose-dependent relationships between cooperating genes in tumorigenesis, a mechanism that has been identified as critical for the function of fusion-oncogenes[13-15]. We foresee this new cell model and vector system being used to study a multitude of potential cooperating oncogenic gene pairs, which could provide mechanistic insights and identify novel therapeutic targets.

## Data Availability Statement

Annotated sequence files are included as FASTA formats in the supplement. Plasmid constructs are in the process of being deposited to Addgene. The data generated in this article are available from the corresponding author by request.

## Acknowledgements

We thank Dave Dunway and the Nationwide Children’s Hospital Flow Cytometry Core for providing services to support this study.

## Author Contributions

Conceptualization: M.K., G.K.; Methodology: M.K., G.K.; Validation: M.K., G.K.; Formal analysis: M.K., A.J., G.K.; Investigation: M.K., A.J., G.K.; Resources: G.K.; Data Curation: M.C., G.K.; Writing – original draft: M.K., G.K.; Writing – review and editing: M.K., A.J., G.K.; Visualization: M.K., G.K.; Supervision: G.K.; Project administration: G.K.; Funding acquisition: G.K.

