## Supplemental Material for "New Dual Inducible Cellular Model to Investigate Temporal Control of Oncogenic Cooperating Genes"

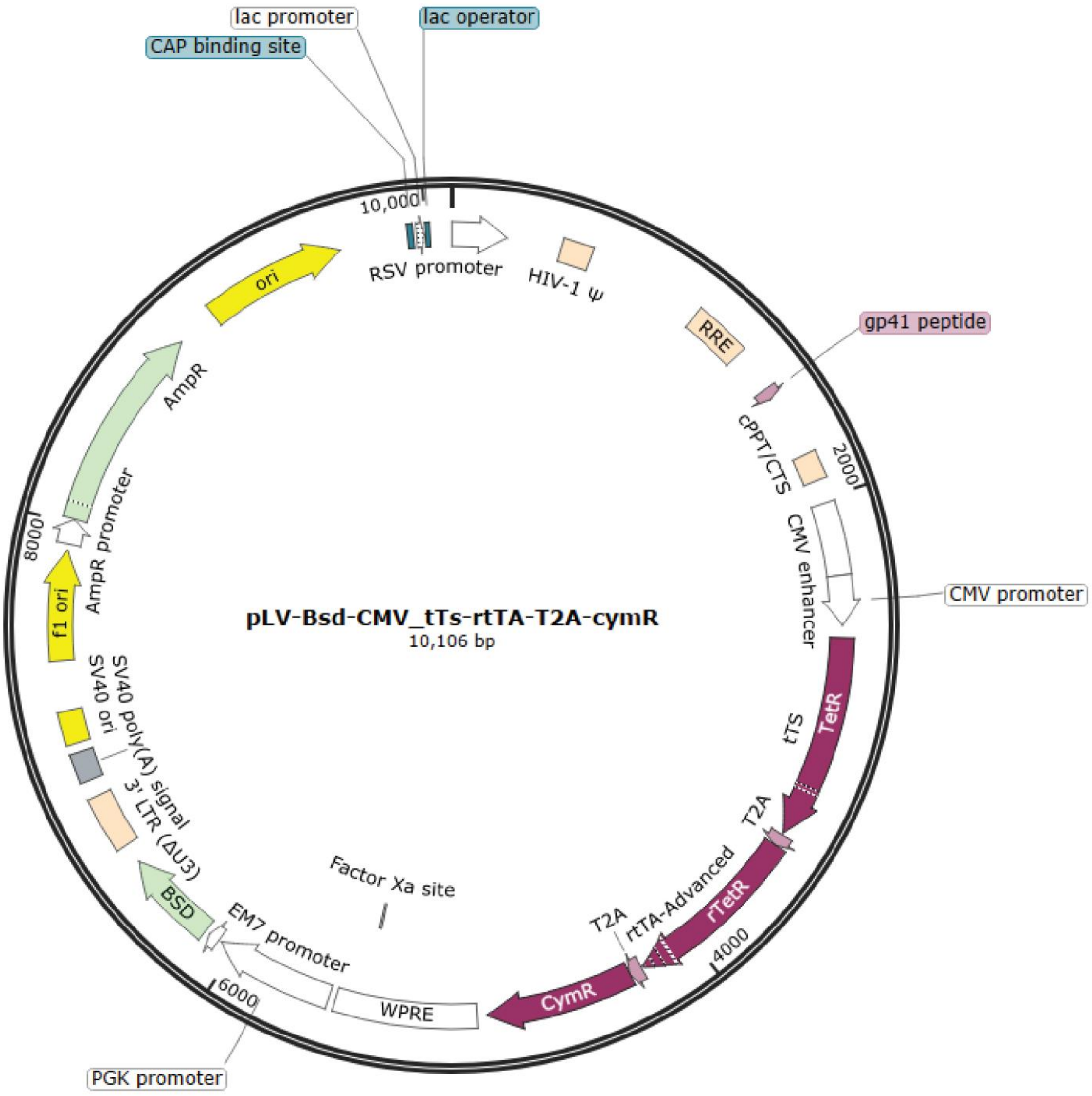


**Figure S1. Lentiviral plasmid map of reverse tet transactivator and cumate repressor.** This lentiviral vector contains the components for the reverse tet transactivator and the cumate repressor (CymR), separated by Thosea asigna virus (T2A) sequences, and all under control of the CMV enhancer/promoter. It also contains a blasticidin resistance gene under control of a phosphoglycerate kinase (PGK) promoter. The sequence can be found in the attached FASTA file.

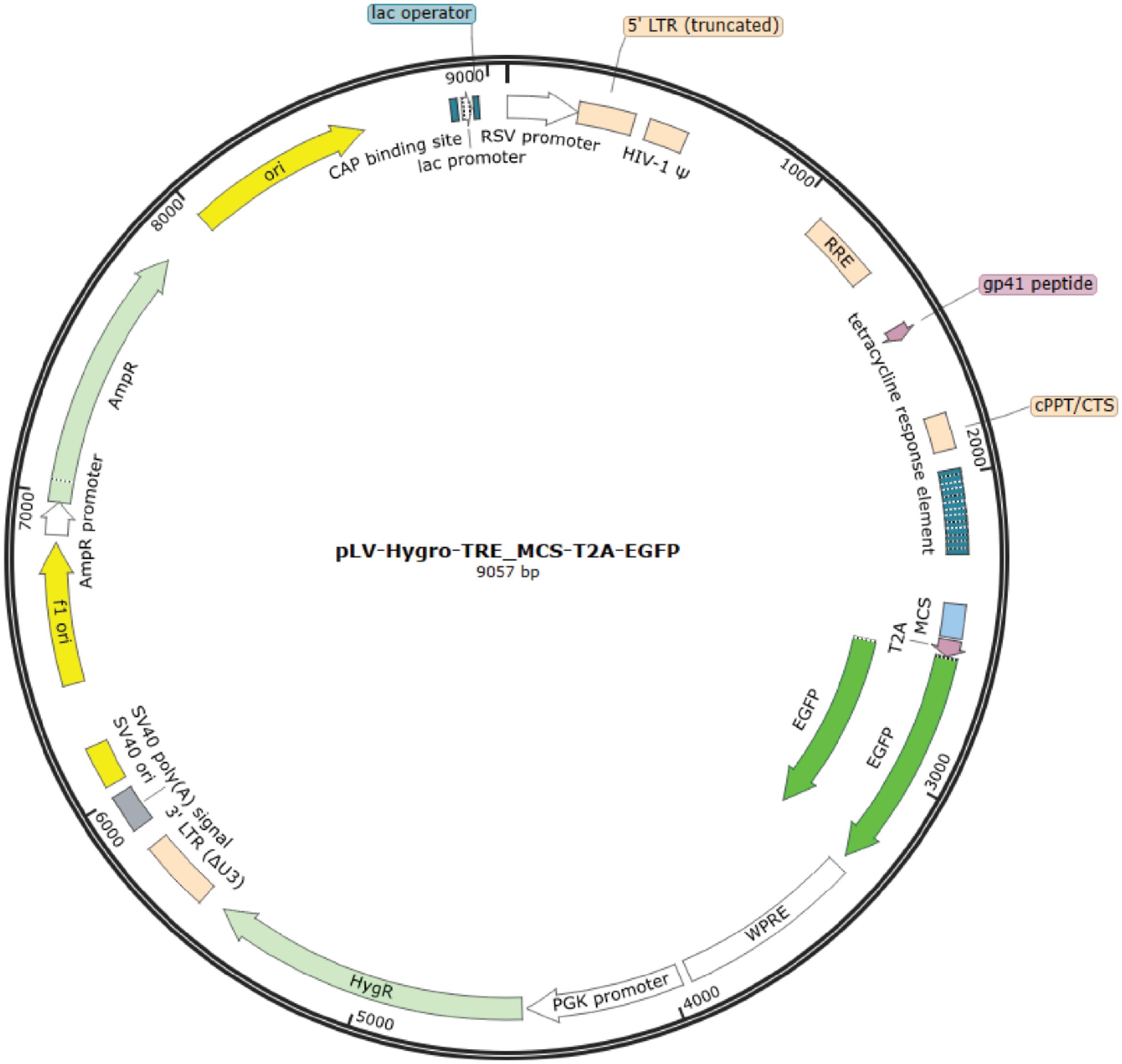


**Figure S2. Lentiviral plasmid map of multicloning site and EGFP.** This lentiviral vector contains a multicloning site and EGFP, separated by a T2A sequence, and under control of the tetracycline response element. It also contains a hygromycin resistance gene under control of a PGK promoter. This sequence can be found in the attached FASTA file.

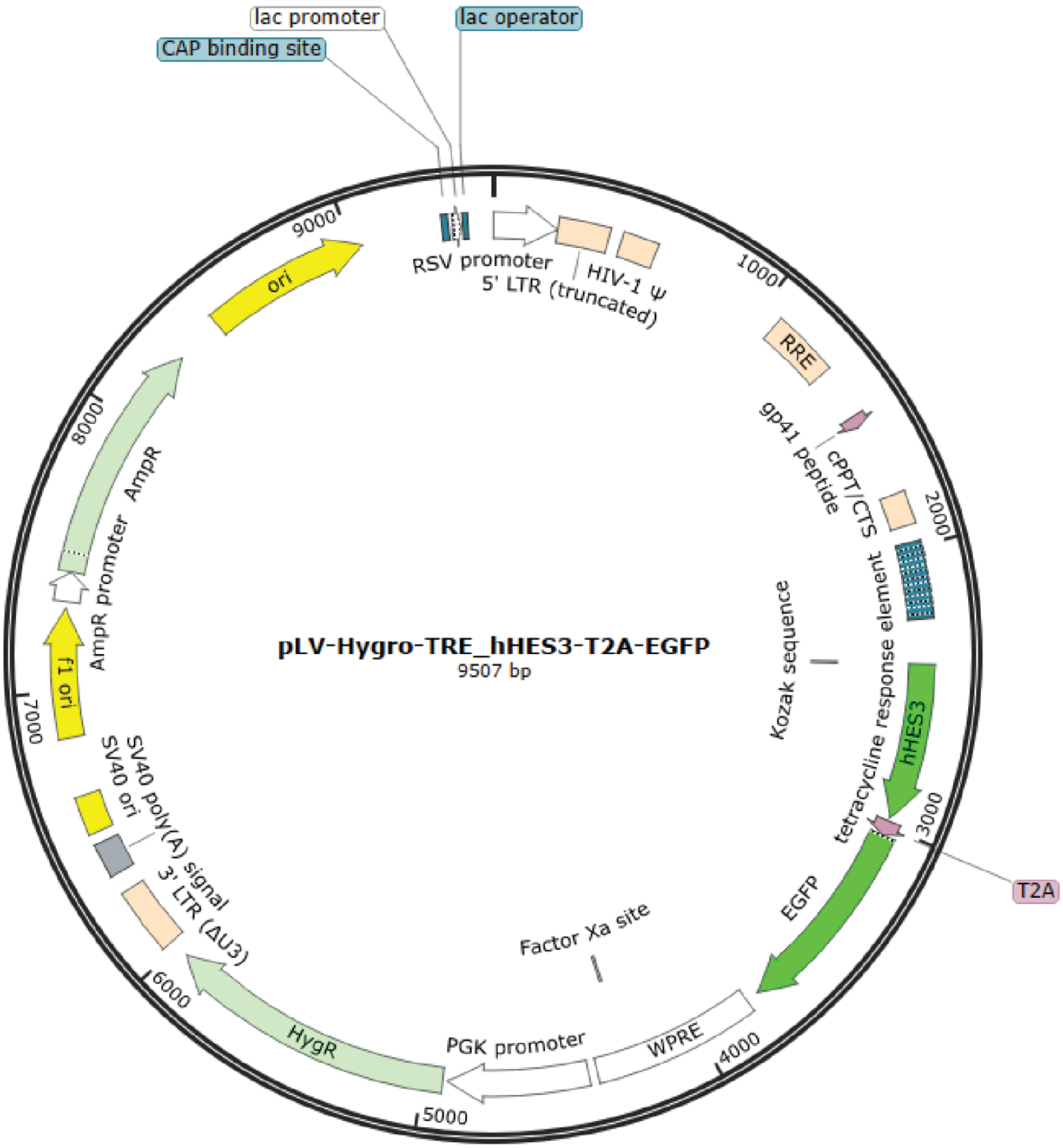


**Figure S3. Lentiviral plasmid map of HES3 and EGFP.** This lentiviral vector contains HES3 and EGFP, separated by a T2A sequence, and under control of the tetracycline response element. It also contains a hygromycin resistance gene under control of a PGK promoter. The sequence can be found in the attached FASTA file.

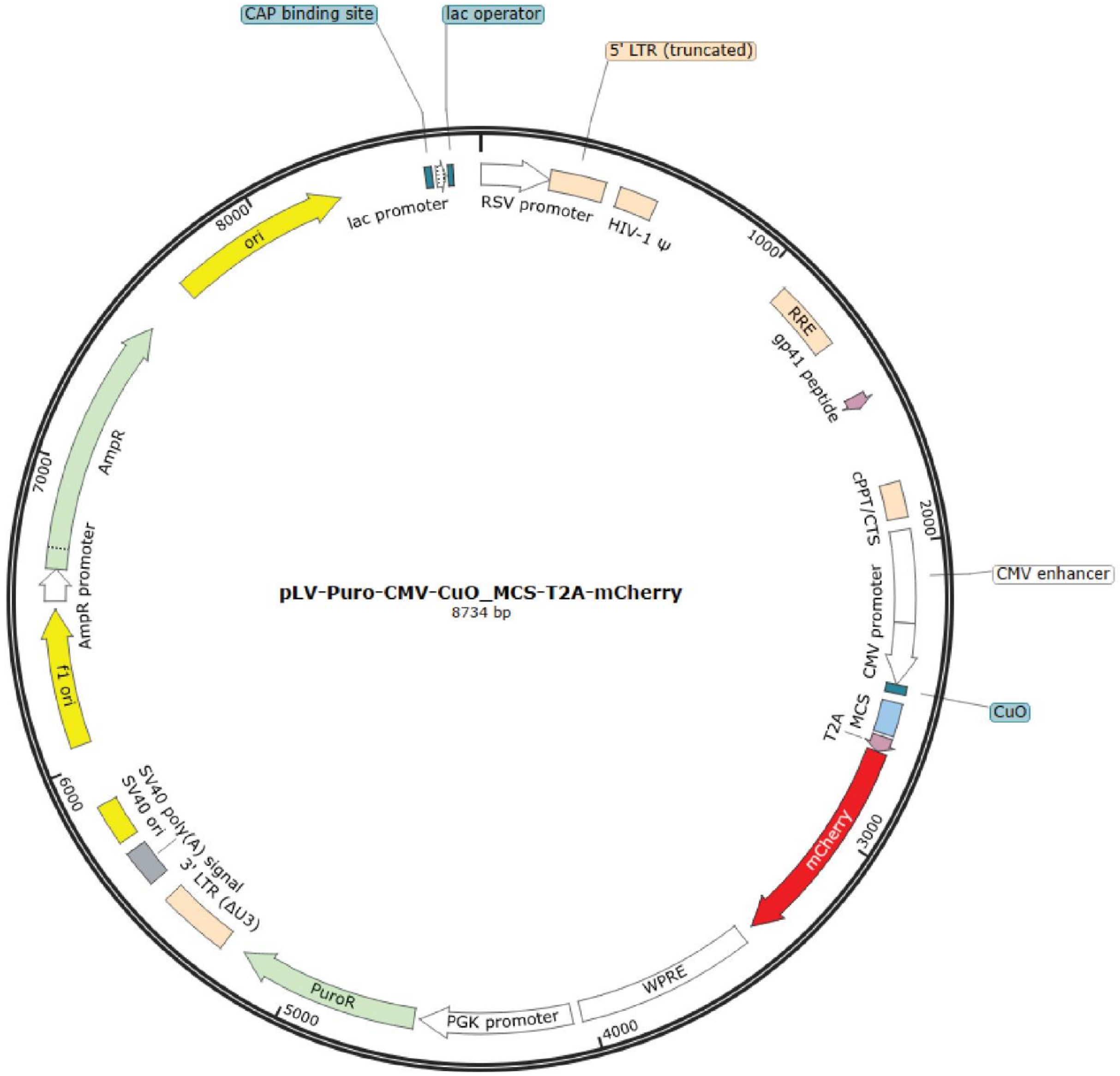


**Figure S4. Lentiviral plasmid map of multicloning site and mCherry.** This lentiviral vector contains a multicloning site and mCherry, separated by a T2A sequence, and under control of the CMV enhancer/promoter with the cumate operator. It also contains a puromycin resistance gene under control of a PGK promoter. The sequence can be found in the attached FASTA file.

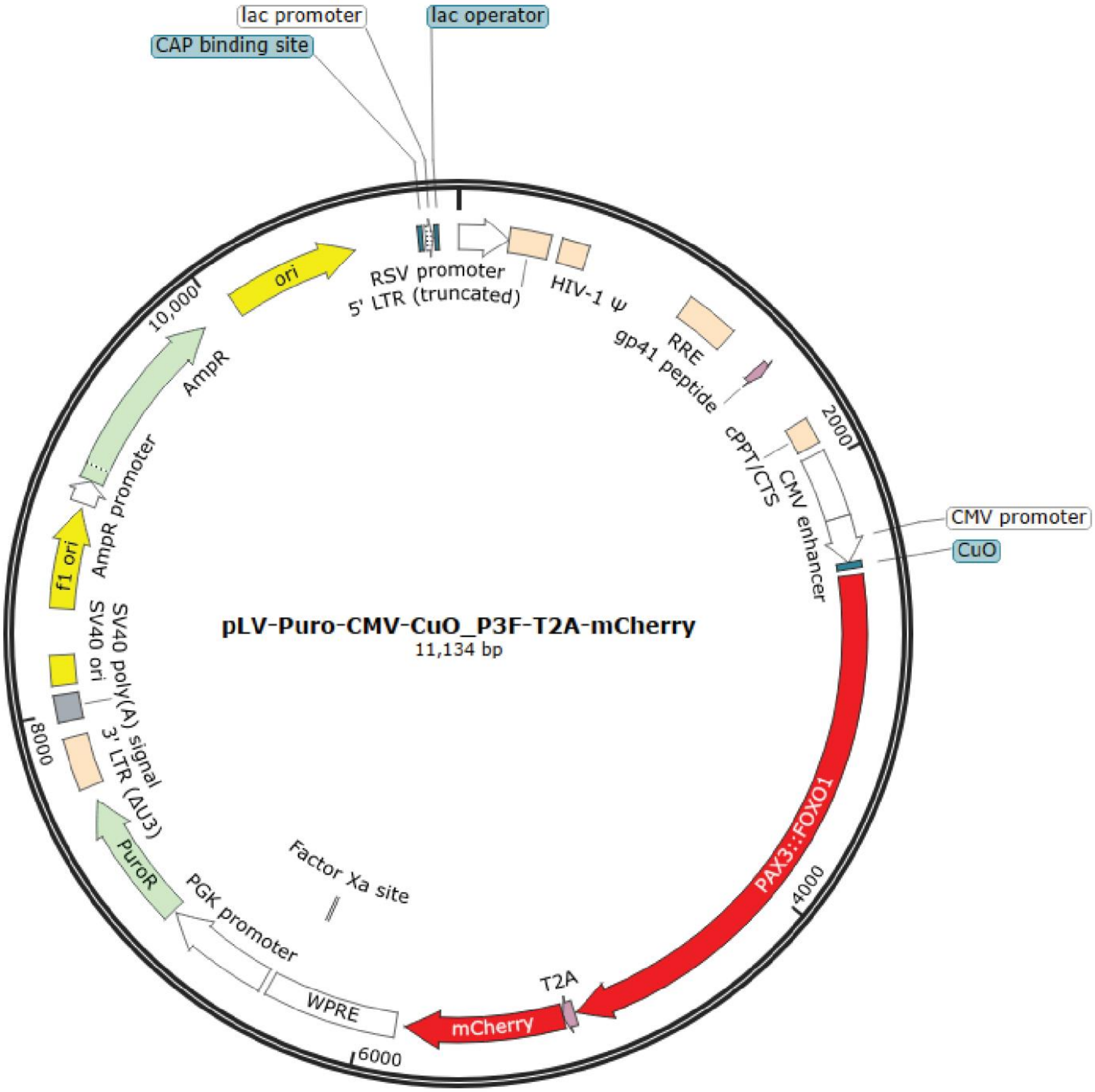


**Figure S5. Lentiviral plasmid map of PAX3::FOXO1 and mCherry.** This lentiviral vector contains PAX3::FOXO1 and mCherry, separated by a T2A sequence, and under control of the CMV enhancer/promoter with the cumate operator. It also contains a puromycin resistance gene under control of a PGK promoter. The sequence can be found in the attached FASTA file.

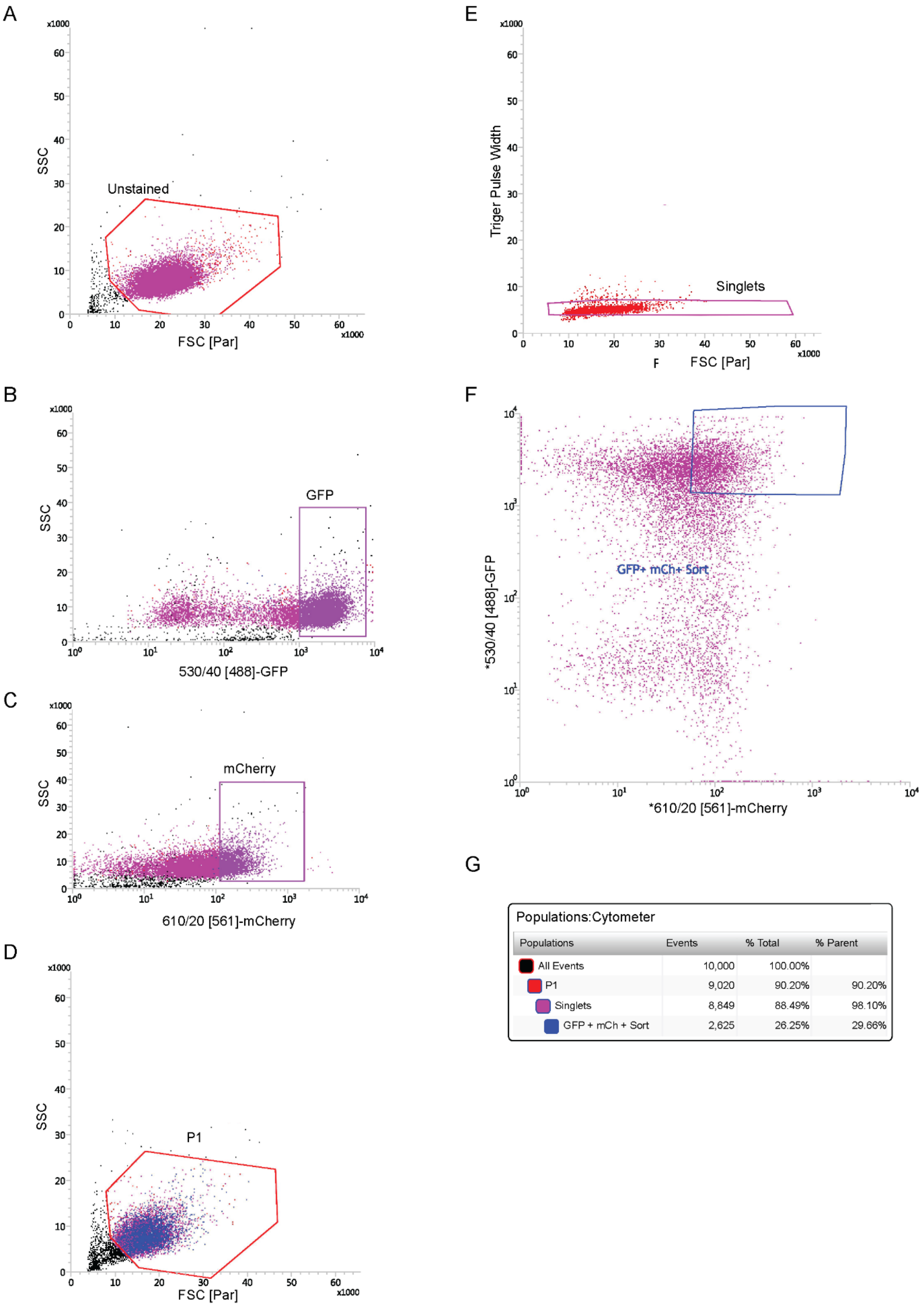


Populations:Cytometer

| Populations | Events | % Total | % Parent |
|---|---|---|---|
| All Events | 10,000 | 100.00% | |
| P1 | 9,020 | 90.20% | 90.20% |
| Singlets | 8,849 | 88.49% | 98.10% |
| GFP + mCh + Sort | 2,625 | 26.25% | 29.66% |

**Figure S6. Fluorescence-Activated Cell Sorting of double-induced cells.** 293Tii cells were seeded on two 6-well plates at 250,000 cells per well. Additionally, naïve 293T cells were seeded on a 6-well plate at 250,000 cells per well. After overnight adherence, anhydrotetracycline was added to three wells of 293Tii cells at 50ng/ml. Cumate was added to three wells of 293Tii cells at 50µg/ml. To the remaining six wells of 293Tii cells, both anhydrotetracycline and cumate were added at 50ng/ml and 50µg/ml, respectively. 24 hours later, additional anhydrotetracycline and cumate were added to the same wells as previous. 24 hours later, cells were collected and diluted to approximately 1,000,000 cells/ml in growth media to be sorted. Naïve 293T cells were used as a negative fluorescence control (**A**). Cells that only received anhydrotetracycline were used to set the gate for GFP fluorescence (**B**). Cells that only received cumate were used to set the gate for mCherry fluorescence (**C**). After setting the two fluorescent gates, double-induced cells were filtered for live cells (**D**). Of the live double-induced cells, singlets were selected for (**E**). Then, live singlets were then sorted into fresh cell growth media, collecting only cells that were both GFP and mCherry double positive (**F**). The final cytometer report of the sort is shown (**G**).
